# IRF8 deficiency causes anxiety-like behavior in a sex-dependent manner

**DOI:** 10.1101/2025.09.02.673764

**Authors:** Stella Zhu, Keita Saeki, Rose-Marie Karlsson, Daniel Abebe, Keiko Ozato

## Abstract

Anxiety disorder is a serious psychiatric disease that affects women twice more than men and disrupts patients’ daily lives. It is often comorbid with major depression and other mental diseases. Various underlying mechanisms have been proposed, such as neurotransmitters and neuroanatomical disruptions, and more recently, oxidative stress; however, much remained unclear, including the role of glial cells. Here, we investigated the role of IRF8 in anxiety disorders in the mouse model. IRF8 is a transcription factor expressed primarily in microglia in the brain. A battery of behavioral tests revealed that female IRF8 knockout (IRF8KO) mice show increased anxiety relative to male IRF8KO and wild-type mice. Female IRF8KO mice also exhibited a higher tendency for obsessive-compulsive disorder. However, these behavioral abnormalities were not observed when IRF8 was deleted postnatally, indicating that it acts during the fetal stage to control anxiety. Transcriptome analysis revealed that IRF8 deficiency leads to redox dysregulation. Further, 2’,7’-dichlorofluorescin diacetate (DCFDA) staining for microglia demonstrated that female IRF8KO microglia produce higher levels of reactive oxygen species (ROS) compared to WT and male IRF8KO counterparts. Detailed RNA-seq analysis, however, did not reveal specific genes that cause high ROS production in female cells. In sum, this work demonstrates that IRF8 in microglia plays a major role in controlling anxiety in a sex dependent manner.

## Introduction

In the United States, nearly 20% of all adults experienced an anxiety disorder in the past year^1, 2^. More than 30% of all adults suffer from some form of anxiety disorder in their lifetime, with women being twice more susceptible than men^2^. Anxiety disorder is characterized as a mental condition that causes persistent, uncontrollable worry and fear, and often panic, that interferes with daily functioning^1^. Anxiety disorders often occur together with major depressive disorder and obsessive-compulsive disorder and can lead to suicidal ideation^3, 4^.

Anxiety disorders occur due to many factors, such as genetics, environmental factors, and temperament^1^. The high comorbidity and overlapping symptoms between anxiety disorders and other psychiatric disorders make isolating genetic risk factors difficult and render therapeutic interventions more complex^5^. Much of the existing research on anxiety disorders has focused on characterizing the neuroendocrine^6, 7^, neurotransmitter^8, 9^, and neuroanatomical disruptions in the limbic system^10, 11^, the emotional-processing brain structures^12, 13^. Increased oxidative stress and inflammation have also been linked to anxiety and depression^14, 15, 16^.

Recent studies indicate that, in addition to neuronal cells, glial cells, particularly microglia, play a pivotal role in anxiety^17, 18^. Microglia, originating from the embryonic yolk sac, are resident immune cells that provide protection against infection in the central nervous system (CNS)^19, 20^. Microglia have ramified processes and monitor the entire brain space for various types of injuries and engage in repair^19, 20^. In addition, microglia have non-immune functions to control the formation and trimming of neuronal synapses^21, 22, 23, 24^. Thus, microglia dysfunction is increasingly implicated in almost all diseases and injuries of the CNS^22, 25^ ^26, 27^. In addition, it is reported that some of the microglia activities are different between males and females^18, 28^.

Several key transcription factors set microglia’s gene expression and functions^29^. IRF8 is a transcription factor expressed primarily in microglia and a few CNS macrophages, such as border-associated macrophages. IRF8 binds to microglia enhancers along with PU.1, Sall1, and directs the development of embryonic and postnatal microglia^30, 31^. IRF8 directs the expression of many microglia identity genes, such as surface receptors, and genes affecting motility and waste clearances^30^. In the periphery, IRF8 promotes the development of bone marrow in myeloid progenitor cells and macrophages to elicit innate immune functions^32, 33, 34^. IRF8 is also involved in osteoclast development^35, 36^.

Here, we show that IRF8 deficiency drives anxiety-like and repetitive disorders in mice in a sex-dependent manner, with female mice exhibiting more severe symptoms. Examination of conditional knock out mice revealed that IRF8 confers protection against anxiety during the prenatal stage. We further observed that female IRF8KO microglia have elevated ROS levels relative to wild type cells. Together, this work identifies microglial IRF8 to inhibit the development of anxiety-like disorders by controlling oxidative stress in a sex dependent manner.

## Methods

### Animals

All experiments described in this study have been performed on mice. They were housed in groups of 2-5 animals under a 12-hour light-dark cycle with ad libitum access to water and food. Experiments were performed with age and sex-matched animals, as stated in the results and figures. Assays were performed on test naïve animals, and pharmacological treatments were administered to drug-naive animals. Constitutive IRF8KO mice (B6(Cg)-Irf8^tm1.2Hm^/J) were described^37^. For conditional deletion of *Irf8* gene, C57BL/6-Irf8^tm1.1Hm^/J (Irf8^fl/fl^), B6.129P2(C)-Cx3cr1^tm2.1(cre/ERT2)Jung^/J (Cx3cr1cre^ERT2^), Rosa26^LSL-YFP^ (R26-EYFP), and control wild-type C57BL/6J (WT) mice were maintained in the NICHD animal facility. Irf8^fl/fl^Cx3cr1^CreERT2/+^R26-EYFP and Irf8^fl/fl^R26-EYFP were crossed to generate Tamoxifen-inducible conditional knockout microglia (IRF8cKO), with a Cx3cr1cre^ERT2^ gene maintained as a heterozygote. The IRF8cKO mice at post-natal day 12 were given 300µg/body Tamoxifen in 50µl Corn Oil intraperitoneally for five consecutive days, and their behavioral phenotype was characterized eight weeks after injection. All animal experiments were performed according to the animal study (ASP#17-044, 20-044, and 23-044) approved by the Animal Care and Use Committees of NICHD, NIH.

### Behavioral tests

Prior to all tests, mice were acclimated to the behavior room for 1 hour, and all experiments were performed between 10:00 am and 3:00 pm with 55-60% room humidity and between 20-23°C. All equipment was cleaned with 1% sodium hypochlorite alcohol before and after each trial to remove odor cues. Unless stated otherwise, the overall activity for each test was recorded using an overhead digital video system and ANY-maze software (Stoelting Co., Wood Dale, IL). When tests were performed in a battery, they were organized from least stressful to most stressful to minimize the effect stress exposure can have on compromising behavior in later tests.

#### Open Field Test (OFT)^38^

The open field test characterizes general locomotor activity and anxiety-like behaviors. Mice were placed in a square box (42×42×42cm) composed of opaque, plastic walls for 60 minutes daily over four days or for 20 minutes at 30 lux background light. This test challenges the rodent’s natural tendencies to explore novel environments and their fear of open, exposed spaces. The time spent in a 14×14cm area in the center (14cm from the walls) was recorded as an index of anxiety-like behavior.

#### Elevated Plus Maze test (EPM)^39^

The maze test combines the mouse’s natural preference for dark spaces and anxiety towards illuminated, open, and elevated areas. The EPM consists of two opposing enclosed arms (40cm each) and two non-enclosed (open) arms shaped as a “plus sign”, and elevated 42cm above the ground. Mice were allowed to explore the platform for 5 minutes at 100 lux background light, and the proportion of time spent in the closed arms was recorded as an index of anxiety-like behavior.

#### Light-Dark Box test (LDB)^40^

The light-dark box test combines the mouse’s natural preference for dark spaces and anxiety towards illuminated, open areas. Mice were placed in a square box (42×42×42cm) with a brightly lit, open compartment with 100 lux background light and an enclosed dark compartment for 10 minutes. The proportion of time spent in the dark compartment was recorded as an index of anxiety-like behavior.

#### Novelty Suppressed Feeding test (NSF)^41^

The novelty-suppressed feeding test assesses how mice reconcile the conflict between an anxiety-inducing context and an appetitive stimulus. Mice were food-restricted for 16 hours and placed in a brightly lit field (42×42×42cm) with 100 lux background light, with a pellet of food in the center. The latency to feed (seconds) was visually noted by an observer as an index of anxiety-like behavior.

#### Nestle Shredding Test (NST)^42^

This test measures repetitive behavior as shredding occurs spontaneously in mice. Mice were placed in a clean cage with fresh bedding and a new nestlet pad for 30 minutes. Nestlet pads were weighed before and after, and the percentage of nestlet shredded was recorded as an index of repetitive behavior.

#### Marble Burying Test (MBT)^42^

This test measures repetitive behavior as burying occurs spontaneously in mice.

Mice were placed in a fresh cage with deep bedding and 20 marbles placed evenly apart (5×4) for 30 minutes. The number of marbles buried, which is manually scored as 2/3 of the marbles being buried below the bedding, is recorded as an index of anxiety.

### Analysis of RNA-seq data

Microglial RNA-Seq data were obtained from P9 and 3-month-old female WT (n=2-4) and IRF8KO (n=2-4) which were deposited previously (GSE231406 and GSE266424)^30^. For male microglia datasets, 15,000 microglia from each mouse were analyzed as described previously (n = 3)^30^. After removing batch effects by sva^43^, EdgeR with the trimmed mean of M values normalization method was used to identify differentially expressed genes (DEGs)^44^. Morpheus (https://software.broadinstitute.org/morpheus/) was used to depict heatmaps, and Metascape was employed for gene ontology analysis^45^.

### Microglia preparation and flow cytometry

All procedures were performed as previously described^30^. In brief, mice were euthanized by CO_2_ asphyxiation and transcardially perfused with 10mL of ice-cold 1x PBS. The brain was harvested onto ice-cold HBSS, chopped, and transferred to a 7ml Dounce and homogenized with quick strokes. The homogenate was then filtered through a 70 μm cell strainer, resuspended in 30% isotonic Percoll, and spun down at 800xg for 30 minutes at 4°C without brake. The intermediate myelin debris was then removed, and the cell pellet was stained with CD11b-BV421 (clone: M1/70, Biolegend), CD45-APC (clone: 30-F11, Invitrogen) and Ly6C-APC/Cy7 (clone: HK1.4, Biolegend) antibodies for 20 min. Sample analysis and cell sorting were performed on FACS AriaIIIµ (BD). 2’,7’-dichlorodihydrofluorescein diacetate (DCFDA; ab113851, Abcam) staining was performed according to manufacturer instructions to measure reactive oxygen species^46^. Fifteen thousand microglia were sorted in BSA-containing ice-cold PBS and stained for 30 min with 20µM DCFDA solution on ice and measured directly by flow cytometry. Data was reanalyzed using FlowJo v10.10.0 software (Tree Star).

## Statistical analysis

All statistical analysis for behavioral tests was performed using R v4.3. To compare the data from four groups (WT females, IRF8KO females, WT males, IRF8KO males), a one-way ANOVA and the Tukey HSD post-hoc test between groups were conducted. A multivariate analysis regressing out genotype, sex, and batch effects was performed to analyze the significance of each factor on the behavior. Drawing all graphs and the unpaired Student’s t-tests for flow cytometry data were carried out using GraphPad Prism 5 software, and a p-value less than 0.05 was considered statistically significant for all cases.

## Data availability

All high-throughput sequence datasets generated in this paper are available in GSE307015 (URLs: https://www.ncbi.nlm.nih.gov/geo/query/acc.cgi?acc= GSE307015). Some omics data used in this paper were obtained from other publications as denoted individually.

## Results

### Female IRF8KO mice exhibit anxiety-like behavior: Initial evidence

Mice lacking IRF8, although having some defects in the myeloid lineage, reach adulthood, breed, and live further in a conventional animal facility. In light of reports that microglia influence behavior in rodent models, we asked whether IRF8KO mice exhibit behavioral abnormalities.^17, 18^. To begin, we carried out a four-day open field test (OFT) for wild type (WT) and IRF8KO mice (Fig.1A)^47^. The OFT brings in environmental fear in mice, which triggers aversive behavior^48^. Through repetitive testing, the mice move less on the next day, and due to habituation, they increase their movement after that^47,49^.

**Figure 1.**
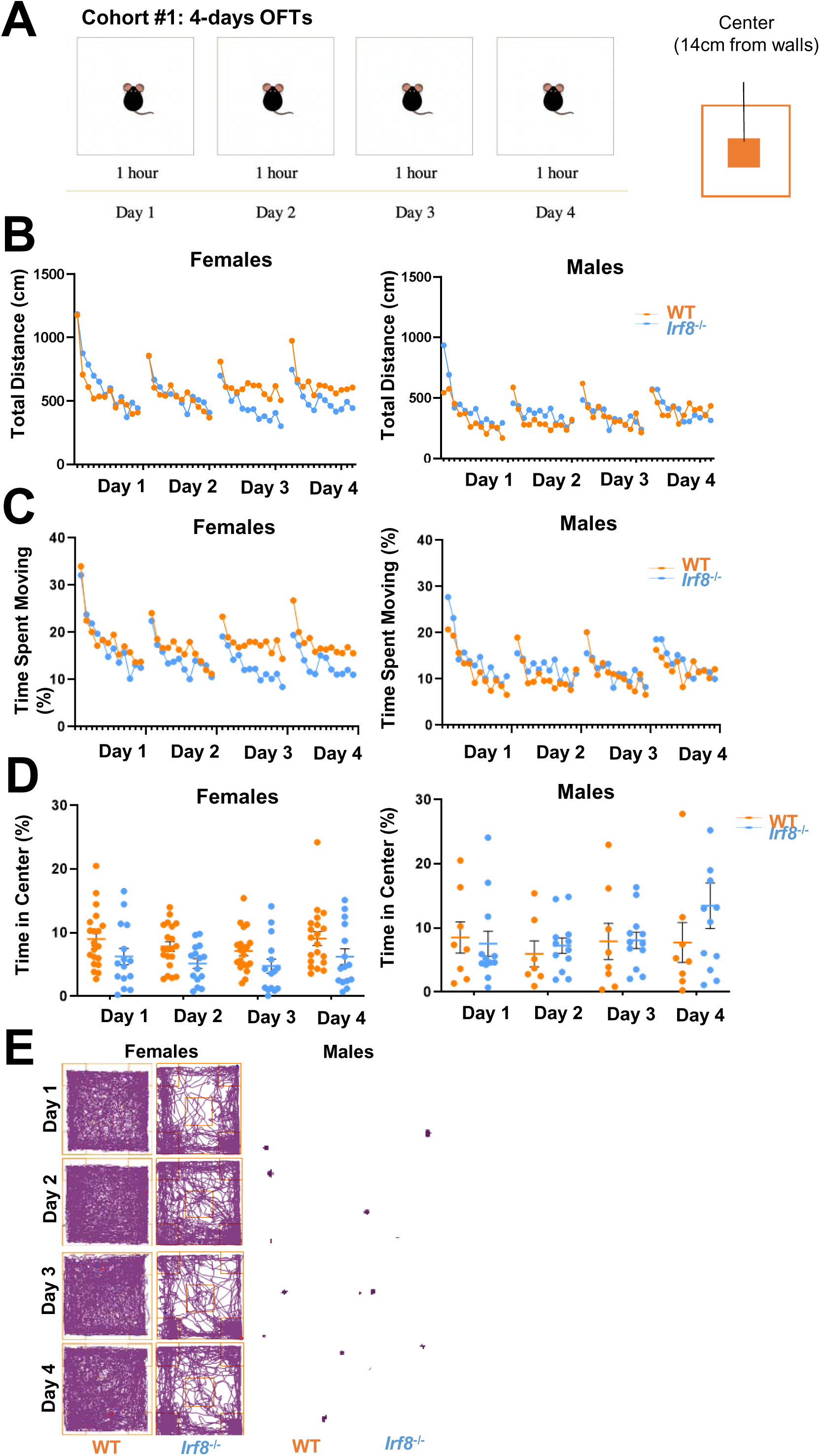
Female IRF8KO mice exhibit anxiety-like behavior: Initial Open Field screening test. **A.** Open field test (OFT) schedule for test-naïve, age-matched (8–11-week-old) WT (n=20 females, n=8 males) and IRF8KO (n=15 females, n=12 males) mice. Mice were placed in the open field for one hour for four consecutive days. **B.** Total distance traveled (cm) on each observed day. Each plot represents the average distance WT (orange) and IRF8KO (blue) mice moved in the arena every 5 minutes. **C.** Proportion of time spent moving on each day. Each plot represents the average of every 5 minutes. **D.** Proportion of time spent in the center area. The mean and standard error bar for each group was also presented. **E.** Representative tracks of individual mice.

This allowed us to assess mouse behavior beyond just movement^47^. Initial video tracking showed that WT and IRF8KO mice move similarly, indicating no locomotor disabilities in IRF8KO mice. Later, however, female IRF8KO mice traveled less than female WT mice (Fig.1B); also, the proportion of time moving was less in female IRF8 mice as they spent more time freezing in the arena, particularly after day 3, compared to female WT mice. Whereas this pattern was not observed in male IRF8KO mice (Fig.1B, C). Next, we analyzed how the mice entered the center of the arena from the periphery. Mice tend to avoid the center of an open field due to fear, allowing us to further examine anxiety-like behavior. Analysis of OFT data revealed that the duration of stay in the center was significantly shorter for female IRF8KO mice than female WT mice, whereas there was no difference between male IRF8KO and male WT mice (Fig.1D), which was also evident by the tracking record (Fig.1E). These results provided a first line of evidence that IRF8 has a role in controlling anxiety-related behavior in a sex-dependent manner.

### Additional tests confirm anxiety-like behavior in female IRF8KO mice

To further probe anxiety-like behavior of IRF8KO mice, we leveraged a battery of additional tests, 20 min OFT, elevated plus maze (EPM), light-dark box (LDB), and novelty suppressed feeding (NSF) ^38^ (diagrams in Fig.2A).

**Figure 2.**
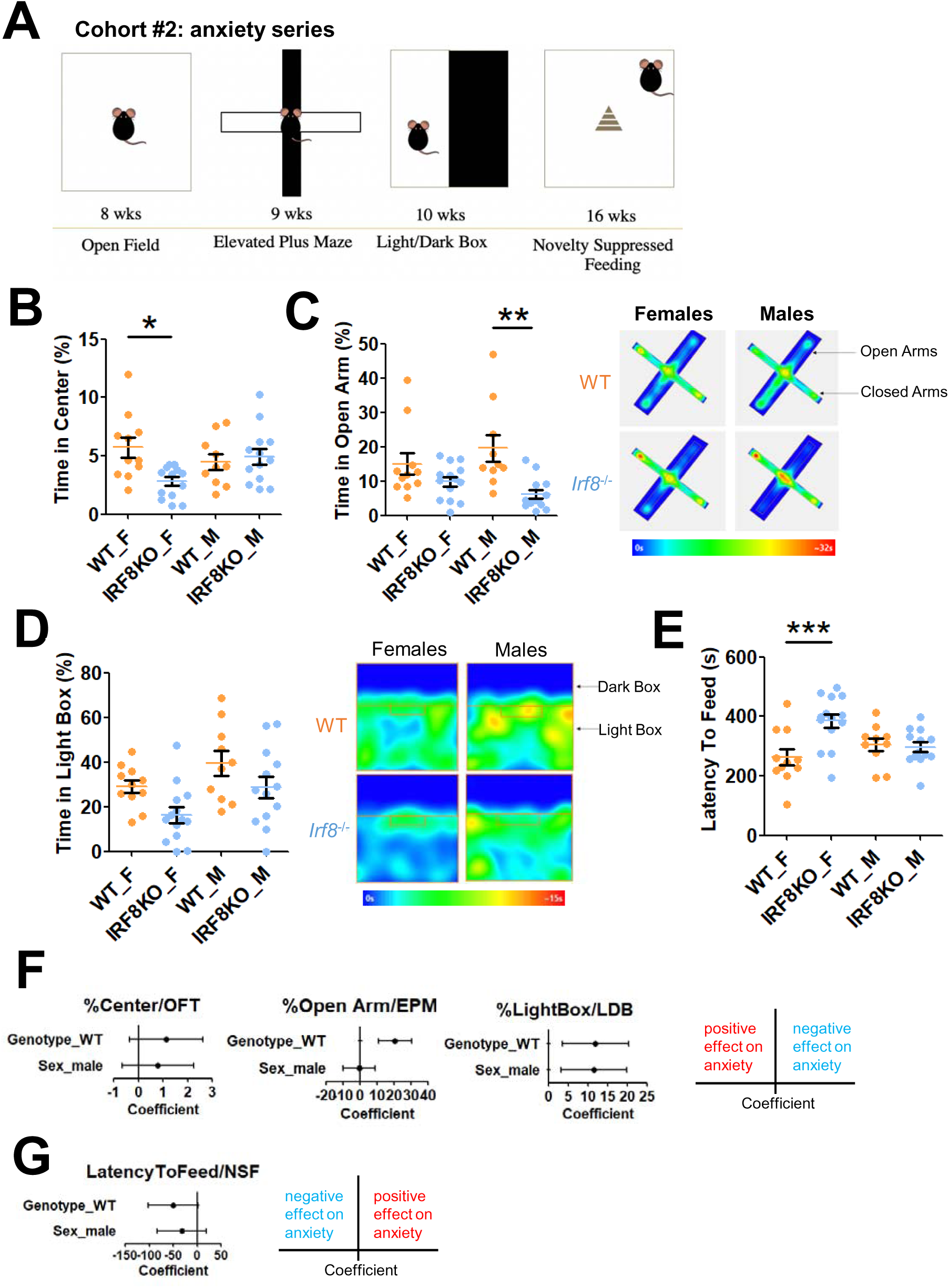
IRF8KO mice displayed anxiety-like behavior, with a more severe phenotype in females: additional tests **A.** Anxiety test battery schedule for test-naïve, age-matched WT (n=11 females, n=14 males) and IRF8KO (n=10 females, n=13 males) mice. We started with 8-week-old mice and performed OFT, EPM, and LDB every week. Then, we waited 6 weeks to minimize the carry-over effects on NSF. **B.** Proportion of time (%) WT (orange) and *Irf8* KO (blue) mice spent moving (left) and they spent in the center area (right) in the Open Field Test. The region 14cm apart from the walls was set as the center area. **C.** Proportion of time spent in the open arms (left) of the elevated plus maze (EPM). Heatmap of averaged group track plots (right). **D.** Time (s) spent in the light box (left) of the light-dark box (LDB). Heatmap of averaged group track plots (right). ANOVA indicated that there is no significant difference in the number of light box entries (F(3, 44)=0.73, p=0.539) and the number of dark box entries (F(3, 44)=2.325, p=0.0878; Fig.S1C). **E.** Latency to feed (seconds) in the novelty-suppressed feeding (NSF). Bar graphs show mean and SEM; Orange (WT) and blue (IRF8KO) nice. Each dot represents an individual data point. Asterisks indicate significant differences (*p.adjusted<0.05, **p.adjusted<0.01, ***p.adjusted<0.001). **F.** Forest plots showing the coefficients of genotype and sex for the specified test, along with the corresponding 95% confidence intervals. A positive value indicates that WT or male mice contribute to higher values in each test, suggesting that they are less susceptible to anxiety stress triggered by the OFT, EPM, and LDB tests. On the other hand, for NSF, a negative value represents milder anxiety because the higher value stands for milder anxiety.

These tests are based on the approach-avoidant paradigm, in which subjects are presented with a threatening environmental stimulus, and their latency to approach or time spent with a potential threat is used as a putative indicator of anxiety^50, 51^. This paradigm is a common strategy to assess anxiety-like behavior in mice. The tests selected here rely on rodents’ natural preference for dark spaces and an aversion toward illuminated, open, and/or elevated areas, as they present an increased risk of predation^51^.

In the first test in the series, a 20 min OFT, the ANOVA indicated that there is a significant difference in time spent in the center among female, male IRF8KO and WT mice (F(3,44)=3.984, p=0.0135). Post-hoc multiple comparison tests revealed that female IRF8KO mice spent significantly less time (p.adjusted=0.0110) in the center than female WT mice, while there was no significant difference in the male IRF8KO and WT mice (p.adjusted=0.959; Fig.2B). There was no significant difference in distance traveled (F(3,44)=2.031, p=0.123; Fig.S1A), In the EPM, there was a significant difference in the number of closed arm entries (F(3,44)=3.08, p=0.0371; Fig.S1B) and the time spent in the open arms (F(3,44)=5.639, p=0.00233; Fig.2C). Post-hoc tests showed that male IRF8KO mice spent significantly less time in the open arm than male WT mice (p.adjusted=0.00230; Fig.2C). Whereas female IRF8KO mice did not show a statistically significant difference from female WT mice (p.adjusted=0.425; Fig. 2C, left). In accordance, the tracks depicting total movement show that IRF8KO mice moved and stayed in the open arms less than WT mice (Fig.2C, right). These results suggest that not only female IRF8KO, but male IRF8KO mice also exhibit increased anxiety-like behavior in this context.

In the LDB assay, there was a significant difference in the time spent in the light box, in that IRF8KO mice spent more time in the Dark relative to WT mice; the tendency was higher in females than males (F(3,44)=5.087, P=0.00413; Fig.2D, left). Total space occupancy corroborated reduced light box entry by IRF8KO mice, more noticeable by the female (Fig.2D, right).

Finally, in the NSF, the ANOVA test revealed a significant difference (F(3, 44)=5.67, p=0.00226) in latency to feed. Accordingly, post-hoc tests revealed that female IRF8KO mice were more reluctant to feed relative to female WT mice (p.adjusted=0.00189; Fig.2E). There was no delay in male IRF8KO mice relative to WT mice. These results collectively support the idea that female IRF8KO mice have increased anxiety-like behavior compared to their counterparts.

Further, multivariate analyses for each test showed that the loss of IRF8 significantly affected EPM and LDB. Additionally, only LDB demonstrated a significant gender bias (Fig.2F). There were also considerable connections between IRF8 loss and OFT and NSF (Fig. 2F), indicating that the combined influence of genotype and sex could contextually impact mouse behavior, resulting in a somewhat diverse phenotype for each test.

### Female IRF8KO mice exhibit compulsive and repetitive behavior

In humans, anxiety disorders are often comorbid with other conditions, such as obsessive-compulsive disorder (OCD)^52^. OCD and anxiety disorders share some neurobiological pathways and genetic risks, thereby are often comorbid, affecting the same patients^53, 54, 55^. Given that OCD is also more prevalent among women, we wondered if our IRF8KO mice likewise exhibit OCD. To that end, we leveraged the nestlet shredding (NST) and marble burying tests (MBT; Fig.3A)^42^. Both tests take advantage of how the target compulsive and repetitive behaviors, shredding and digging, occur spontaneously in natural rodent activity^42^. We found that female IRF8KO mice showed increased NST activity compared to female WT mice, while male IRF8KO and WT mice showed lower NST activity (Fig.3B). Accordingly, the ANOVA found that there was a significance (F(3,60)=4.378, p=0.00748). Additionally, post-hoc tests revealed that female IRF8KO mice shredded a significantly greater percentage of the nestlet (p.adjusted=0.00346) compared to their WT counterpart, whereas there was no remarkable difference in shredding activity between male WT and male IRF8KO mice (p.adjusted=0.9866; Fig.3B). Photographic record also verified increased shredding activity by female IRF8KO mice (Fig.3B, right).

**Figure 3.**
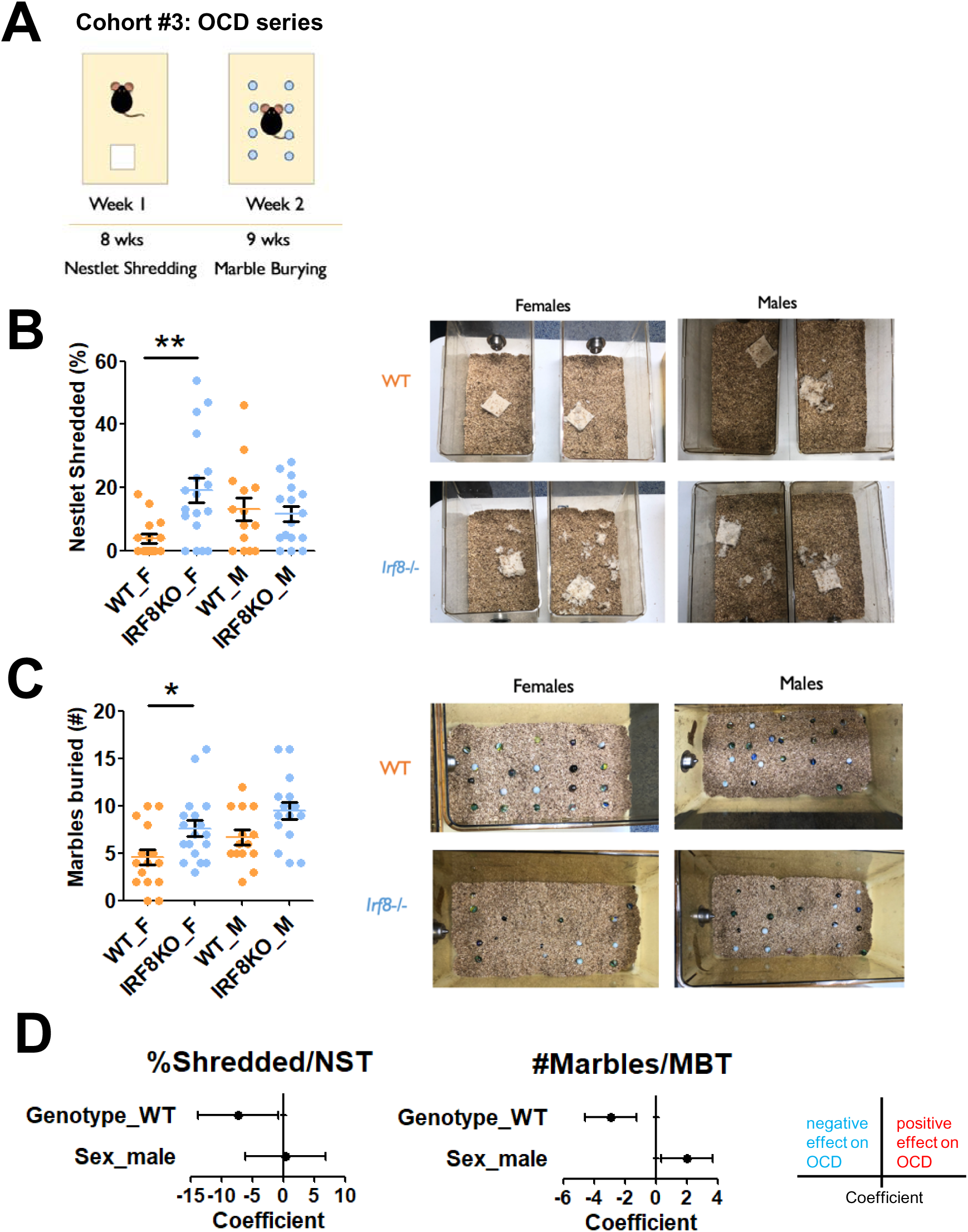
IRF8KO mice exhibit repetitive and compulsive behavior, with a more severe phenotype in females. **A.** Test schedule for naïve, age-matched WT (n=16 females, n=14 males) and IRF8KO (n=18 females, n=16 males) mice. The age of mice when each test was conducted was indicated in the scheme. **B.** Percentage of nestlet shredded (left) during the nestlet shredding test (NST). Representative pictures (right) of nestlets shredded. **C.** Numbers of marbles buried (left) during the marble burying test (MBT). Representative pictures (right) of marbles buried. Bar graphs show mean and SEM; Orange (WT) and blue (IRF8KO) circles are individual data points. Asterisks indicate significant differences (*p.adjusted<0.05, **p.adjusted<0.01). **D.** Forest plots showing the coefficients of genotype and sex on the indicated test, along with 95% confidence intervals. A positive value suggests that WT or male mice contribute to the obsessive behavior.

In the MBT, the ANOVA first verified that there was a significant difference (F(3, 60)=6.023, p=0.00118) in the number of marbles buried. In line with this, post-hoc comparisons revealed that female IRF8KO mice buried significantly more marbles than WT mice (p.adjusted=0.04929; Fig.3C). Photographic records also support the results in the post-hoc tests (Fig.3C, right). However, male IRF8KO mice, although they appeared to bury somewhat more marbles than male WT mice, the difference was not statistically significant (Fig.3C). Multivariate analyses showed that both genotype and sex independently impacted the number of buried marbles, while only genotype significantly affected the NST (Fig. 3D). Together, both sexes of IRF8KO mice had higher OCD tendency than WT mice. WT mice, however, had greater sexual dichotomy than IRF8KO mice. These data suggest that anxiety disorders and OCD have shared as well as distinct mechanisms.

### Postnatal deletion of IRF8 does not cause anxiety-like behavior

IRF8 is expressed in microglia throughout life. It is required for the embryonic development of microglia^56^. In addition, IRF8 is required for postnatal, functional maturation of microglia^30, 56^. It has been shown that neural circuits are influenced by gender at around E18^57, 58^, and those linked to anxiety behavior continue to mature until around postnatal day 60^59, 60^. To investigate at which stage IRF8 affects anxiety-related behavior, we examined conditional IRF8 knockout (IRF8cKO) mice, where IRF8 was deleted postnatally upon Tamoxifen injection^30^. Deletion of IRF8 was confirmed before the behavioral test for each mouse^30^. Female IRF8cKO and WT mice were examined for all the behavioral tests above, OFT, EMT, and LDB for anxiety, and NST plus MBT for OCD (Fig. 4A diagram). To our surprise, IRF8cKO mice did not display anxiety-like behavior, nor OCD, in that scores were similar between IRF8cKO and WT mice for each test (Fig. 4B-F). These results indicate that IRF8 sets neuronal mechanisms and pathways that prevent anxiety and OCD during the prenatal stage.

**Figure 4.**
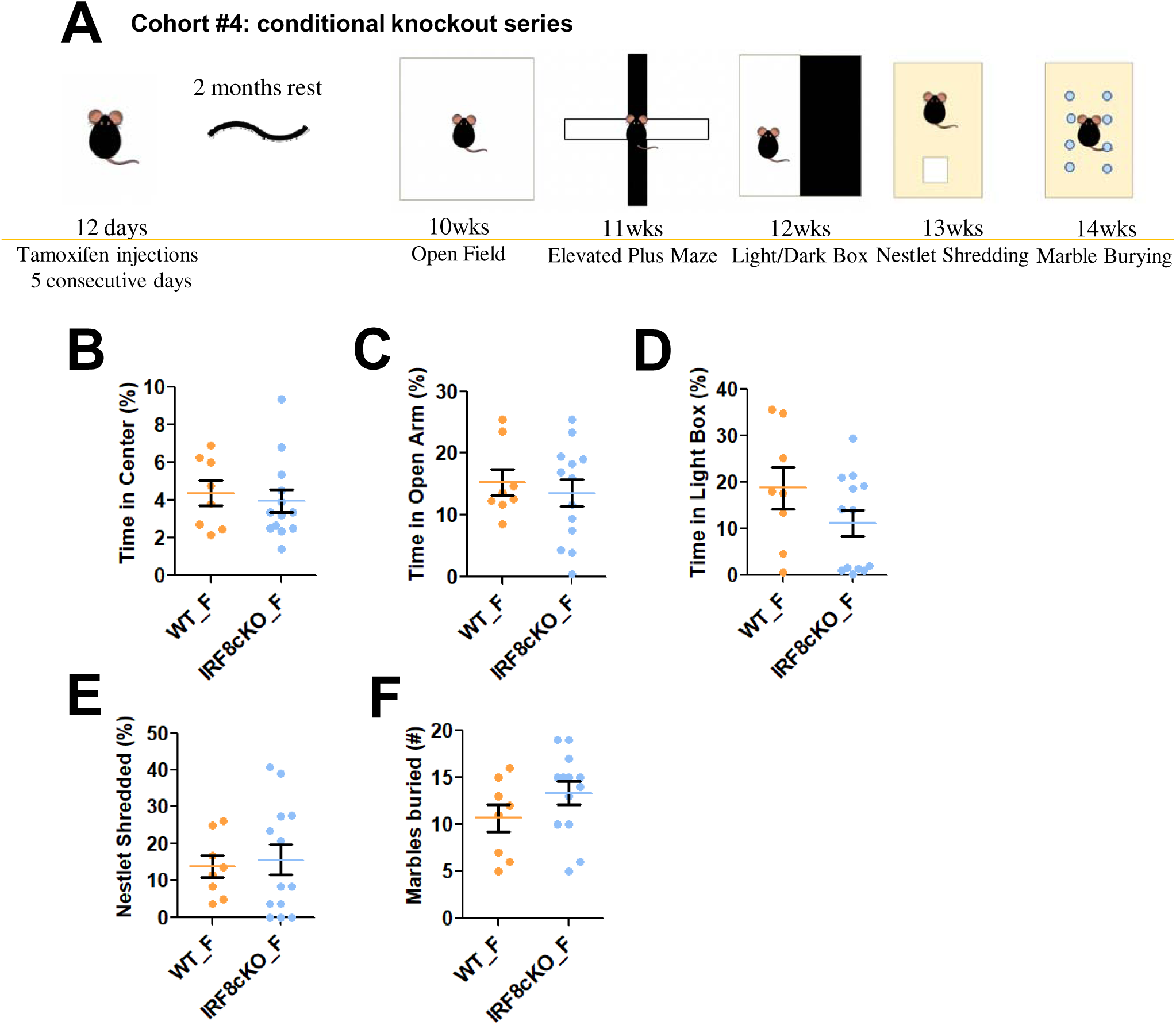
Postnatal deletion of IRF8 does not cause anxiety-like behavior **A.** Anxiety and repetitive test battery schedule for test-naïve, age-matched female WT (n=8) and IRF8cKO (n=12) mice. IRF8floxCx3cr1CreERT2 mice of 12 days old age were given Tamoxifen for five consecutive days to delete *Irf8* gene specifically in microglia. The age of mice when each test was conducted was indicated in the scheme. **B.** Proportion of time spent moving of the 20 minutes OFT (See also Fig. 2B). **C.** Time spent in the open arms of the elevated plus maze (EPM; See also Fig. 2C). **D.** Time spent in the light of the light-dark box (LDB; See also Fig. 2D). **E.** Percentage of nestlet shredded during the nestlet shredding test (NST; See also Fig. 3B). **F.** Numbers of marbles buried (left) during the marble burying test (MBT; See also Fig. 3C). Orange (WT) and blue (IRF8cKO) circles are individual data points. Bar graphs show mean and SEM.

### IRF8 regulates ROS production in microglia in a sex dependent manner

Previous reports indicated that oxidative stress is causally linked to anxiety and depression^61, 62, 63^. We first examined our RNA-seq data comparing WT and IRF8KO microglia (Experimental schemes in Fig.5A)^30^. GO analysis pointed to oxidative phosphorylation as DEGs (Fig.5B, S2A). In accordance, we found an increase in genes associated with oxidative phosphorylation and mitochondria (Fig.5C). Particularly, IRF8KO microglia expressed higher levels of mitochondrial Complex I and Complex IV genes, which include *Ndufa1* and *Cox4i1* (Fig.5C; S2B for males). Mitochondrial Complex I comprises 14 core protein complexes and 30 accessory subunits, with the latter being crucial for Complex I assembly and function^64^. In adult IRF8KO microglia, 12 out of 30 accessory proteins (40%) were upregulated over WT microglia. In addition, IRF8KO microglia from postnatal day 9 showed higher expression of Complex I genes compared to the WT counterparts (Fig.S2C). Additionally, almost all genes for the Complex IV proteins were expressed at higher levels in adult and postnatal day 9 IRF8KO cells than WT cells, which was more evident in P9 microglia (Fig. S2C). These data possibly suggest that IRF8KO microglia have higher oxidative phosphorylation.

**Figure 5.**
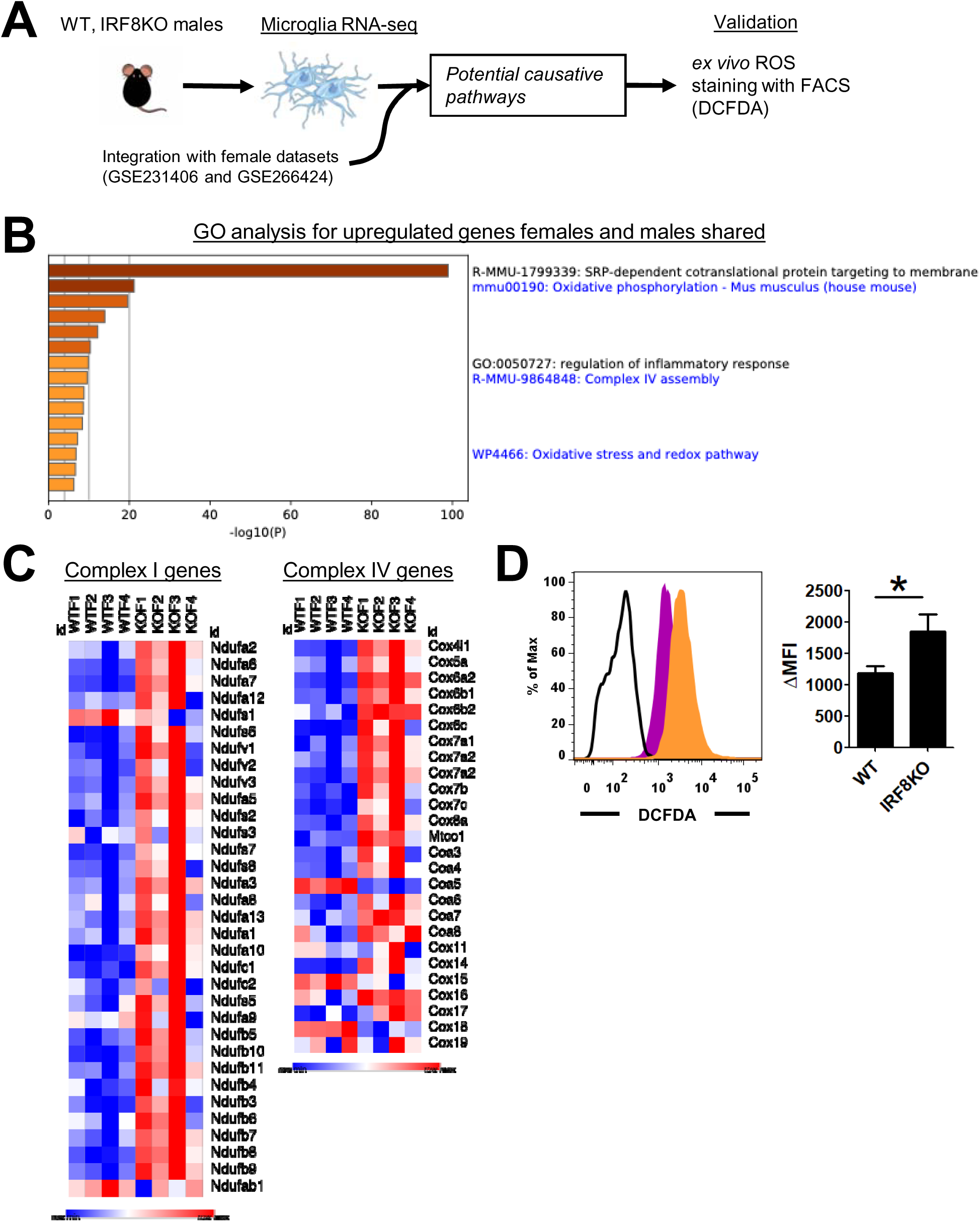
IRF8 regulates ROS production in microglia in a sex-dependent manner **A.** Workflow for analyzing microglia from WT and IRF8KO mice. The deposited RNA-seq data (GSE231406; adult WT and IRF8KO microglia datasets, n=4 females and GSE266424; P9 and adult WT and IRF8KO female microglia datasets, n=2-4) were reanalyzed with the new datasets of male microglia (n=3). Then, ex vivo microglia were stained with DCFDA dye and analyzed by FACS. **B.** GO analysis for the DEGs expressed higher in IRF8KO microglia, regardless of sex. The shared genes were identified in Fig.S2A. Terms of interest were selected from the Metascape output. **C.** Heatmaps for mitochondrial Complex I (NAD(P)H+ dehydrogenase; left) and Complex IV (cytochrome c oxidase; right) gene expression in adult, female IRF8KO and WT microglia. **D.** Flow cytometry of sorted microglia stained with a DCFDA dye. WT (magenta) and IRF8KO (orange) microglia were presented on the left. The mean fluorescent intensity (MFI) was analyzed on the right. Bar graphs show mean and SEM; Asterisks indicate significant differences (*p.adjusted<0.05).

These data are interesting, since the mitochondrial complex is a source of reactive oxygen species (ROS) production^64^.

We next asked whether the amounts of ROS differ between IRF8KO and WT microglia, and whether there is a difference between male and female. Microglia were isolated by FACS sorting under low temperature in order to maintain in vivo transcriptome programs^30^. To measure ROS levels, live microglia were incubated with 2’,7’-dichlorodihydrofluorescein diacetate (DCFDA) at 4^0^C, and ROS were detected in flow cytometry (Fig.5D)^46^. We avoided using other dyes that require incubation at higher temperatures, which would globally alter microglia transcriptomes^65, 66^. Remarkably, gender-wise comparisons revealed that only female IRF8KO microglia showed significantly higher DCFDA signals compared to female WT microglia (Fig. 5D, top left). In contrast, male IRF8KO microglia had lower DCFDA staining, the amounts of which were somewhat lower than those of male WT microglia (Fig. S2D). These data pointed to the possibility that female IRF8KO microglia have higher levels of ROS, perhaps due to less effective redox regulation^67, 68^. We anticipated that comparison of transcriptomes between female and male microglia from IRF8KO mice would identify differentially expressed genes (DEGs) that are relevant to redox regulation, thus providing candidate genes responsible for increased ROS production in female cells. Integrative RNA-seq analysis was performed for female and male microglia from IRF8KO and WT mice (Fig.5A, S2D). To our dismay, DEGs did not include genes for ROS production, nor those involved in regulating ROS levels, despite DEGs were mostly those genes known to be sex-linked (Fig S2D). Therefore, a simple comparison of RNA-seq data from female and male cells did not explain why female IRF8KO microglia produce more ROS. These data indicate that the mechanisms of high ROS production in female IRF8KO microglia are complex and cannot be pinpointed to specific genes (see Discussion).

## Discussion

In this study, we investigated the role of the transcription factor IRF8 in regulating behavioral attributes in mice. We focused on IRF8, since it is expressed predominantly in microglia and a few CNS macrophages in the brain, and it controls microglia gene expression. Using a series of anxiety-avoidant tasks, we found that mice without IRF8 display anxiety disorder-like behavior. These abnormalities are presumably due to unwarranted fear in IRF8KO mice, which discourages exploratory activities. IRF8 deficient mice also exhibited compulsive and repetitive tendencies, in line with the link between anxiety and OCD noted in humans^4, 55^. Particularly, this behavioral phenotype manifested more severely in female IRF8KO mice than in males, revealing the same gender bias observed in human anxiety disorders^69^.

The coupling of anxiety disorder and OCD may indicate that they have a shared mechanism preserved through evolution. Anxiety disorder and OCD apparently have anatomically common routes, in that anxiety disorder is linked to disruption of the amygdala and the prefrontal cortex^12, 70^, while cortical regions are implicated in OCD^71,72^.

How do microglia regulate behavior and what is the role of IRF8? Microglia play a critical role in the development of the brain and other CNS cells^21, 22, 23, 24^. They direct the formation of neuronal synapses and myelination not only during the embryonic stage but throughout life. They scan the entire brain space to detect injuries and engage in repair. Besides, microglia detect incoming pathogens and elicit anti-pathogen resistance. IRF8 is essential for the development of microglia during fetal and early postnatal stages^30, 56^. Many of microglia’s functions in adults also depend on continuous expression of IRF8^30^. We recently reported that IRF8 binds to enhancers for many genes that provide microglia-specific functions and that deletion of IRF8 deprives microglia of their identity and functionality^30^. The absence of IRF8 likely prevents the formation of neuronal connections important for handling stress and anxiety. However, deletion of IRF8 after birth did not cause anxiety-like behavior, nor OCD, indicating that IRF8 sets proper neuronal connections during the prenatal stage to control anxiety.

Our effort to understand mechanisms of anxiety and repetitive behavior in IRF8KO mice led us to compare microglia transcriptome profiles. We found that mitochondria Complex I and IV were upregulated in IRF8KO cells relative to WT cells. Mitochondrial Complex I plays a crucial role in regulating oxidative stress. It’s a major site of ROS production where ROS is continuously generated from the leakage of electrons to form superoxide O ^.-51^. These data pointed out the significance of redox regulation in shaping behavioral traits. Our data also unravels the previously unappreciated role of IRF8 in redox regulation.

A most salient finding in our analysis was that female IRF8 microglia produced higher levels of ROS than female WT and male cells, as evidenced by DCFDA staining. Excessive ROS reacts with lipids, nucleic acids, and proteins, leading to deleterious effects on surrounding cells and tissues. Increased ROS production leads to prolonged oxidative stress, a pathological hallmark of neuroinflammatory and neurodegenerative diseases and aging^73^. It has been amply reported that anxiety-disorder-like behavior is prompted by oxidative stress, which could be mitigated by neutralizing ROS under certain conditions^16, 63, 74^. The increased ROS production in female IRF8KO microglia likely results in prolonged unremittent oxidative stress. The data that female IRF8KO cells have increased ROS paralleled the data that female IRF8KO mice suffered increased anxiety-like behavior, highlighting the shared sexual dichotomy.

Although it might have been anticipated, inspection of DEGs between female and male microglia did not identify redox related genes that could explain increased ROS production in female IRF8KO cells. This may not be unexpected, considering that the ROS levels are determined by many factors^67^. There appear to be more than 100 genes that influence ROS levels. It is possible that these genes exert their effect in combination, rather than alone, when the expression of each is below the DEG cut-off. In addition, the difference may occur at the level of proteins rather than RNA expression. Besides, antioxidants derived from external and internal sources could affect ROS levels in microglia. Moreover, it is possible that other factors, such as sex hormones, exacerbate the anxiety and repetitive behavior of female IRF8KO mice^25^.

In summary, we show that IRF8, a transcription factor expressed in microglia, influences the development of anxiety disorders in a sex dependent manner. Our data point to oxidative stress caused by elevated ROS as one of the potential underlying mechanisms.

## Acknowledgements

This research was supported [in part] by the Intramural Research Program of the National Institutes of Health (NIH). The authors’ contributions are considered Works of the United States Government. The findings and conclusions presented in this paper are those of the authors and do not necessarily reflect the views of the NIH or the U.S. Department of Health and Human Services. We thank A. Dey, J. Kassis, and members of the NICHD DIR affinity group for discussion on behavioral experiments and critical reading of the manuscript.

## Inclusion and ethics statement

All the contributors to this study have met the authorship criteria required by Nature Portfolio journals and have been included as authors because their participation was essential for the design and implementation of the study. Roles and responsibilities were determined among collaborators prior to the research. This work encompasses locally relevant findings that have been identified in collaboration with local partners. The research was not significantly restricted or prohibited in the researchers’ environment and did not lead to stigmatization, incrimination, discrimination, or personal risk to participants. Citations of local and regional research relevant to our study have been considered.

## Author Contributions Statement

SZ and KS performed experiments, analyzed data, and wrote the manuscript; RMK and DA supervised behavior studies; KO supervised the study and wrote the manuscript.

## Supplemental Figure Legend

**Figure S1.**

**A.** Total distance (m) that mice traveled in the OFT arena.

**B.** The number of mouse entries to the closed arm (left) and open arm (right) in EPM test.

**C.** The number of mouse entries to the dark box (left) and light box (right) in LDB test. Bar graphs show mean and SEM; Orange (WT) and blue (IRF8KO) circles are individual data points. Asterisks indicate significant differences (*p.adjusted<0.05, **p.adjusted<0.01, ***p.adjusted<0.001).

**Figure S2.**

**A.** Venn diagrams displaying the shared genes between two types of DEGs. The top diagram compares upregulated DEGs (FDR<0.01) in male IRF8KO microglia with those in female IRF8KO microglia. The bottom diagram compares downregulated DEGs in male IRF8KO microglia with those in females.

**B.** Heatmaps for mitochondrial Complex I (NAD(P)H+ dehydrogenase; left) and Complex IV (cytochrome c oxidase; right) genes expression in adult, female IRF8KO and WT microglia.

**C.** Heatmaps for mitochondrial Complex I (NAD(P)H+ dehydrogenase; left) and Complex IV (cytochrome c oxidase; right) genes expression in P9 and adult female IRF8KO and WT microglia. The datasets deposited in GSE266424 was re-analyzed.

**D.** Flow cytometry of male, adult microglia stained with a DCFDA dye. WT (magenta) and IRF8KO (orange) microglia were presented in the left. The mean fluorescent intensity (MFI) was analyzed in the right.

**E.** Volcano plots presenting the expression of genes that are differentially expressed in females (pink) and males (blue) microglia from WT (left) and IRF8KO (right). Male microglia specifically expressed Y-linked genes, such as *Kdm5d* and *Ddx3y*, while X chromosome genes, such as *Xist* and *Tsix*, were expressed only in females. Note that there was a substantial number of genes irrelevant to sex chromosomes that were upregulated or downregulated in a genotype-specific manner.

