## Supplemental Figures (S1 and S2) for "IRF8 deficiency causes anxiety-like behavior in a sex-dependent manner"

### Slide 1
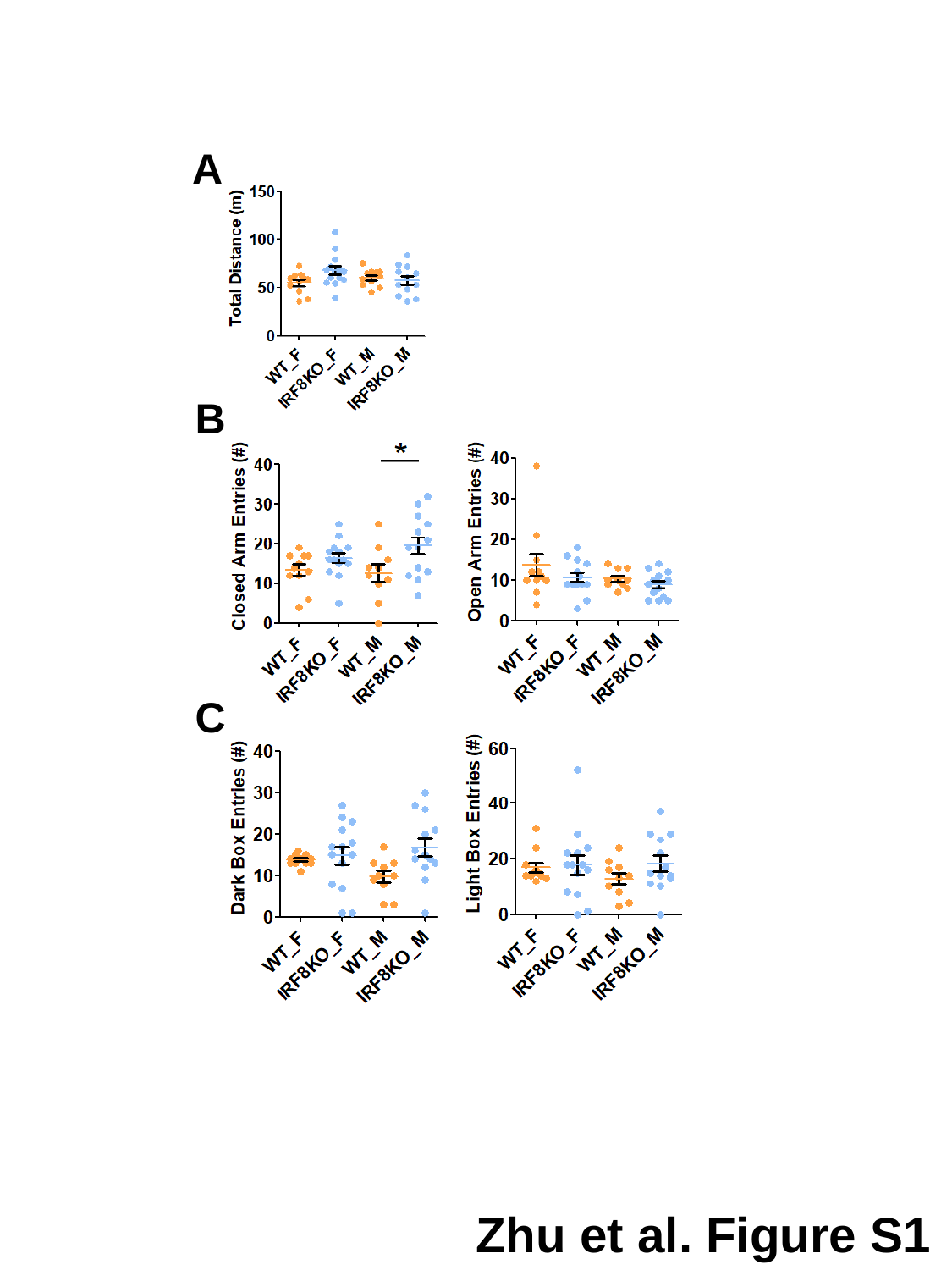

A
B
C
Zhu et al. Figure S1

### Slide 2
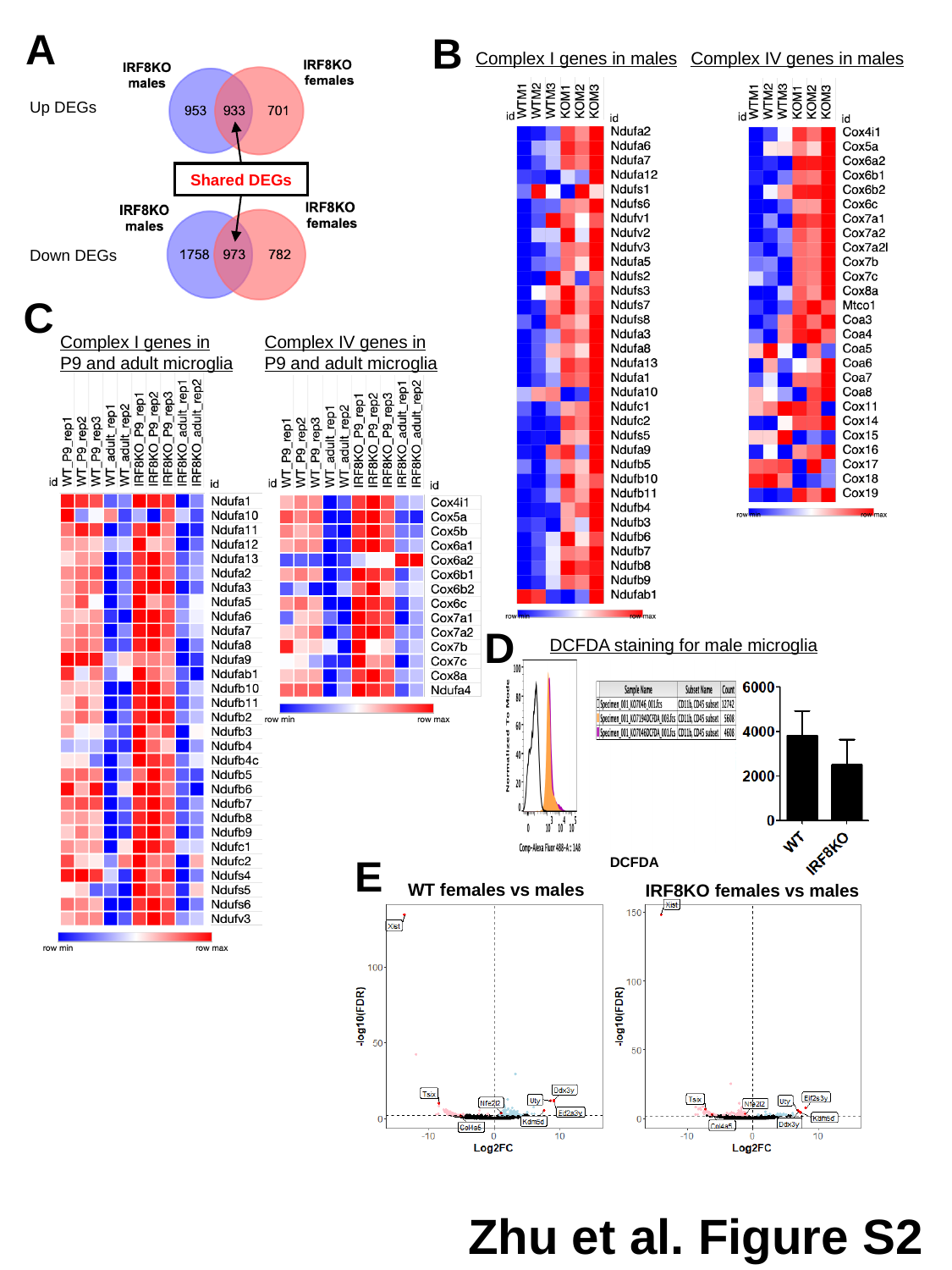

A
B
Complex I genes in males
Complex IV genes in males
Up DEGs
Shared DEGs
Down DEGs
C
Complex I genes in P9 and adult microglia
Complex IV genes in P9 and adult microglia
D
DCFDA staining for male microglia
E
DCFDA
WT females vs males
IRF8KO females vs males
Zhu et al. Figure S2
